# Multi-omics analysis reveals key immunogenic signatures induced by oncolytic Zika virus infection of paediatric brain tumour cells

**DOI:** 10.1101/2024.11.28.625843

**Authors:** Matt Sherwood, Thiago G. Mitsugi, Carolini Kaid, Brandon Coke, Mayana Zatz, Kevin Maringer, Oswaldo K. Okamoto, Rob M. Ewing

## Abstract

Brain tumours disproportionately affect children and are the largest cause of paediatric cancer-related death. Despite decades of research, paediatric standard-of-care therapy still predominantly relies on surgery, radiotherapy, and systemic use of cytotoxic chemotherapeutic agents, all of which can result in debilitating acute and late effects. Novel therapies that engage the immune system, such as oncolytic viruses (OVs), hold great promise and are desperately needed. Zika virus (ZIKV) infects and destroys aggressive cells from paediatric medulloblastoma, atypical teratoid rhabdoid tumour (ATRT), diffuse midline glioma (DMG), ependymoma and neuroblastoma. Despite this, the molecular mechanisms underpinning this therapeutic response are grossly unknown. By profiling the transcriptome across a time-course, we comprehensively investigated the response of paediatric medulloblastoma and ATRT brain tumour cells to ZIKV infection at the transcriptome level for the first time. We observed conserved TNF signalling pathway and cytokine signalling-related signatures following ZIKV infection. We demonstrated that the canonical TNF-alpha signalling pathway is implicated in oncolysis by reducing the viability of ZIKV-infected brain tumour cells and is a likely contributor to the anti-tumoural immune response through TNF-alpha secretion. Our findings have highlighted TNF-alpha as a potential prognostic marker for oncolytic ZIKV virotherapy. Performing a 49-plex ELISA, we generated the most comprehensive ZIKV-infected cancer cell secretome to date. We demonstrated that ZIKV infection induces a clinically relevant and diverse pro-inflammatory brain tumour cell secretome, thus circumventing the need for transgene modification to boost efficacy. We assessed publicly available scRNA-Seq data to model how the ZIKV-induced secretome may (i) interact with medulloblastoma tumour microenvironment (TME) cells via paracrine signalling and (ii) polarise lymph node immune cells via endocrine signalling. Our modelling has provided significant insight into the cytokine response that orchestrates the diverse anti-tumoural immune response during oncolytic ZIKV infection of brain tumours. Our findings have significantly contributed to understanding the molecular mechanisms governing oncolytic ZIKV infection and will help pave the way towards ZIKV-based virotherapy.

## Introduction

Malignant paediatric central nervous system (CNS) tumours, including medulloblastoma and ATRT, are the most common solid childhood cancer and are the leading cause of mortality in paediatric cancer patients (1). Current therapy regimens are aggressive and frequently result in debilitating long-term sequelae that ultimately impair the quality of life of patients who do not succumb to fatal tumour recurrence. There is a clear and unmet need for less toxic and more targeted therapies, especially those capable of co-opting the immune system against the tumour. Oncolytic viruses (OV) selectively infect and kill cancer cells, thus minimising long-term sequelae by reducing the need for high doses of chemotherapy and radiotherapy. Their efficacy stems from a two-pronged mechanism of action where the virus can directly lyse the infected cancer cells (oncolysis) and elicit an anti-tumoural immune response. Many cancers, including medulloblastoma and ATRT, contain cancer stem-like cells that drive poor prognosis factors such as elevated tumour heterogeneity, metastasis and therapeutic resistance (2,3). The ability of OVs to target these aggressive stem-like cells is a unique advantage in overcoming the therapeutic resistance of these highly heterogeneous and malignant cancers. To date, four OVs have been clinically approved, and > 200 OV clinical trials are underway to treat aggressive forms of cancer (4). OV research has gained momentum since 2015, when the first FDA approval was granted for a modified form of herpes simplex virus type 1 (T-VEC) to treat recurrent adult melanoma (5). In 2022, a breakthrough in the fight against brain cancer was made when the Japanese Ministry of Health, Labor and Welfare approved the oncolytic herpes virus Delytact (G47Δ) to treat residual or recurrent adult glioblastoma (6). A dozen clinical trials have assessed OV therapy for paediatric brain tumour OV, half of which are completed. Some trials report improved paediatric cancer patient survival, and crucially, all report that OVs are primarily accompanied by only low-grade adverse events (7–11).

Zika virus (Orthoflavivirus zikaense) is a positive-sense single-stranded RNA virus (+ssRNA) in the *Orthoflavivirus* genus. Maternal ZIKV infection during the first, second and third trimesters of human pregnancy results in vertical transmission in 47%, 28% and 25% of cases, and congenital ZIKV syndrome (CZS) symptoms in 9%, 3% and 1% of cases, respectively (12). ZIKV infects fetal radial glial (RGC) and neural precursor (NPC) cells, leading to cell death and growth reduction in the fetal brain (13–16). Conversely, postnatal ZIKV infection is primarily asymptomatic, with the disease generally being self-limiting and symptoms resolving within two to seven days (17). Both ZIKV-infected children and adults suffer from the same main acute symptoms: headache, fever, rash (exanthema), joint pain (arthralgia), conjunctivitis, and muscle pain (myalgia) (18,19). In rare instances, more severe conditions such as Guillain-Barré Syndrome, meningitis and encephalitis can occur; however, these primarily affect adults rather than children (17,20). Work by ourselves and others has shown that ZIKV can infect and destroy aggressive cells from paediatric (medulloblastoma, ATRT, DMG, ependymoma and neuroblastoma) and adult (glioma and meningioma) nervous system tumours (21–28). We demonstrated that ZIKV effectively targets and destroys human CNS tumour cells and can inhibit metastatic spread without causing neurological damage in xenograft mouse models (21,29). ZIKV-induced tumoural immune cell infiltration has been documented to include lymphoid (CD4+ T, CD8+ T and NK cells) and myeloid (monocytes, macrophages, DCs and microglia) cells (30–32). This leads to ZIKV-induced tumour clearance and long-term immunity against tumour cells in immunocompetent glioma mice models, which is dependent on CD4+ and CD8+ T cells (30–32). In a pre-clinical study, members of our research team confirmed ZIKV efficacy against spontaneous intracranial canine tumours (33). Crucially, ZIKV reduced tumour volume, induced an anti-tumoral immune response, improved clinical symptoms, and did not cause any persisting adverse conditions (33).

Large data and omics techniques have revolutionised our understanding of biological systems. As these techniques are yet to be applied to oncolytic ZIKV infection, the biology governing the response in paediatric brain tumour cells is generally unknown, with WNT signalling being the only known molecular mechanism involved (21). Assessing oncolytic ZIKV infection of tumour cells alongside clinically relevant CZS patient-derived NPCs is important to tease out any common or differential mechanisms that ZIKV utilises for either tumour cell oncolysis or its fetal neuropathology. Despite this, CZS patient-derived NPCs are yet to be incorporated into oncolytic ZIKV research. The efficacy of most OVs is heavily dictated by the ability to induce immunogenic cell death (ICD) and a concomitant anti-tumoural immune response (34). It is currently unclear how intracellular or extracellular factors, such as signalling pathways or cytokine secretione, may contribute to ZIKV ICD and orchestrate the anti-tumoural immune response.

In the current study, we first performed a transcriptomic temporal survey and observed cell type-specific responses of paediatric brain tumour cells and CZS patient-derived NPCs during early ZIKV infection (12-24 hours post-infection (hpi)). NPCs are highly sensitive to ZIKV infection and undergo vast downregulation of neurodevelopmental processes and essential upstream signalling pathways, likely underpinning the development of CZS in these patients (35). In ZIKV-infected paediatric brain tumour cells, we observe transcriptomic signatures indicative of ICD, TNF pathway and cytokine signalling responses. We show the upregulation of canonical and non-canonical TNF signalling pathways following infection, and we identify TNF- alpha as a potential prognostic marker for brain tumour ZIKV OV therapy. Investigating cytokine signalling, we generate the most comprehensive ZIKV-infected cancer cell secretome to date and show that ZIKV induces a diverse and predominantly pro-inflammatory secretome from brain tumour cells. Assessing publicly available scRNA-Seq data, we employ an *in silico* approach to model how the ZIKV secretome may contribute to an anti-tumoural immune response. We achieve this by assessing how our identified ZIKV-induced pro-inflammatory secretome may interact with medulloblastoma TME cells via paracrine signalling or polarise lymph node immune cells via endocrine signalling. In summary, our findings shed light onto the molecular mechanisms underpinning oncolytic ZIKV infection of paediatric brain tumour cells and contribute to the ultimate goal of developing an oncolytic ZIKV virotherapy for paediatric oncology.

## Results

### ZIKV infection induces cell type-specific transcriptome responses

Following ZIKV infection, we observe distinct viral sub-cellular staining from 12 hours post-infection (hpi) in the aggressive paediatric USP7 ATRT and USP13 medulloblastoma cell lines (Figure 1A). Viral titration primarily identifies the release of mature viral particles between 12 hpi and 24 hpi, indicating 12-24 hpi as a biologically relevant timeframe during early ZIKV infection (Figure 1B). We infected USP7 (P7) and USP13 (P13) brain tumour cells and NPC-763-1 (N3) and NPC-788-1 (N7) neural precursor cells for 12, 18, and 24 hours and we performed RNA-Seq. We observed that both in the presence and absence of ZIKV infection the brain tumour cell transcriptomes highly contrast that of the NPCs (Figure 1C, Supplementary Figure 1A). In most instances, hierarchical clustering and Principal Component Analysis (PCA) group the RNA-Seq samples by experimental condition, cell line, and disease state (Figure 1C, Supplementary Figure 1B). Assessing ZIKV genome expression, we observe approximately 32, 131 and 356 times more viral genomes in the tumour cells than in the NPCs at 12, 18 and 24 hpi, respectively (Figure 1D). This highlights a greater propensity for ZIKV to replicate in brain tumour cells than in NPCs, possibly due to the dysregulation of anti-viral responses during transformation. Despite the low accumulation of ZIKV genomes in the NPCs, thousands of differentially expressed genes (DEGs) are observed (Table 1), supporting the well-known highly sensitive nature of NPCs to ZIKV infection.

**Figure 1.**
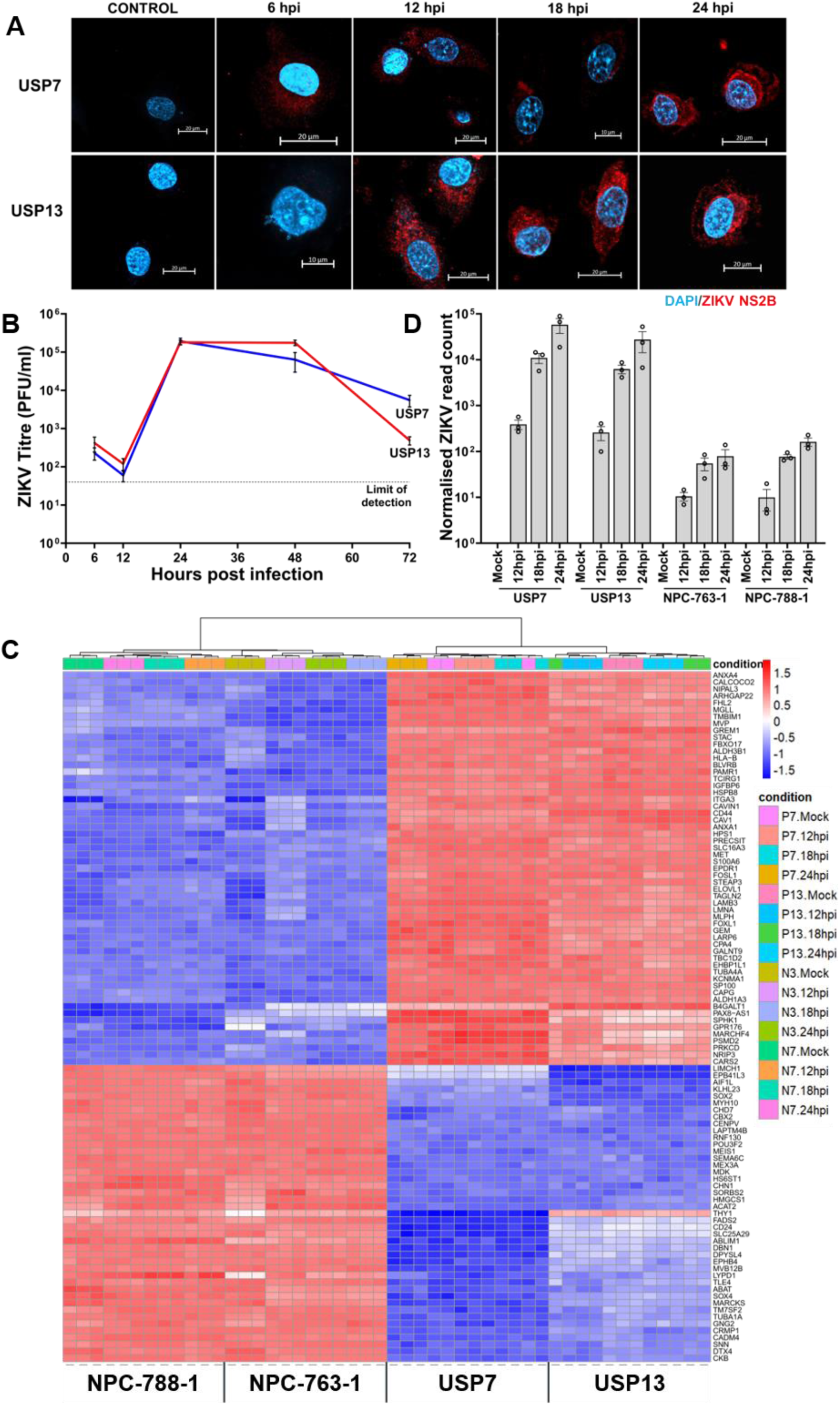
ZIKV infection dynamics in brain tumour and neural precursor cells. **(A)** Confocal microscopy of ZIKV-infected USP7 and USP13 brain tumour cells across the first 24 hours of infection, staining for ZIKV replication (ZIKV NS2B, red) and the nucleus (DAPI, blue). **(B)** ZIKV titre in USP7 and USP13 cells across 72 hours of infection (Mean ± SEM, N=4). **(C)** Heatmap of the Top 100 differential genes across all 48 RNA-Seq samples. Hierarchical clustering arranged the heatmap by condition (columns) and genes (rows); the dendrogram for the latter was omitted to aid visualisation. **(D)** RNA-Seq ZIKV genome count across the different infection conditions in USP7, USP13, NPC-763-1, and NPC-788-1 cells (Mean ± SEM, N=3). Abbreviations, Zika virus (ZIKV), USP7 (P7), USP13 (P13), NPC-763-1 (N3), NPC-788-1 (N7), hours post-infection (hpi), adjusted p-value (padj).

**Table 1.** Table of the differentially expressed genes (DEGs) observed following ZIKV infection in all four cell lines.

|  | Differentially Expressed Genes |  |  |  |
| --- | --- | --- | --- | --- |
|  | P7 | P13 | N3 | N7 |
| **12 hpi** | 446 | 2012 | 3826 | 1726 |
| **18 hpi** | 86 | 325 | 3066 | 2049 |
| **24 hpi** | 2349 | 3017 | 2644 | 1732 |

Gene Ontology (GO) analysis identifies a multitude of differentially regulated biological processes in ZIKV-infected brain tumour and neural precursor cells (Figure 2A). We observe vast downregulation of developmental biological processes in ZIKV-infected NPCs that contribute to the CZS development, including morphogenesis, CNS development, neuron migration, and differentiation (Figure 2A). Multiple developmental signalling pathways which govern and coordinate these biological processes are also differentially regulated in ZIKV-infected NPCs, including WNT, Hippo, MAPK, Notch, TGF-beta, and PI3K-Akt signalling pathways (Figure 2B). The largest differential gene, biological process, and pathway responses in brain tumour cells occur at 24 hpi (Table 1, Figure 2A and 2B). By 24 hpi, multiple terms implicated in cell interaction and the extracellular matrix are downregulated, whilst apoptosis, protein ubiquitination, protein processing in endoplasmic reticulum and negative regulation of transcription from RNA polymerase II promoter terms are upregulated in both ZIKV-infected brain tumour cells (Figure 2A and 2B). Multiple cell cycle and DNA replication/repair terms are significantly upregulated specifically in USP13 cells during 12-18 hpi, whereas cell cycle-related terms are significantly downregulated in USP7 at 24 hpi, indicating that ZIKV may differentially regulate brain tumour cell growth.

**Figure 2.**
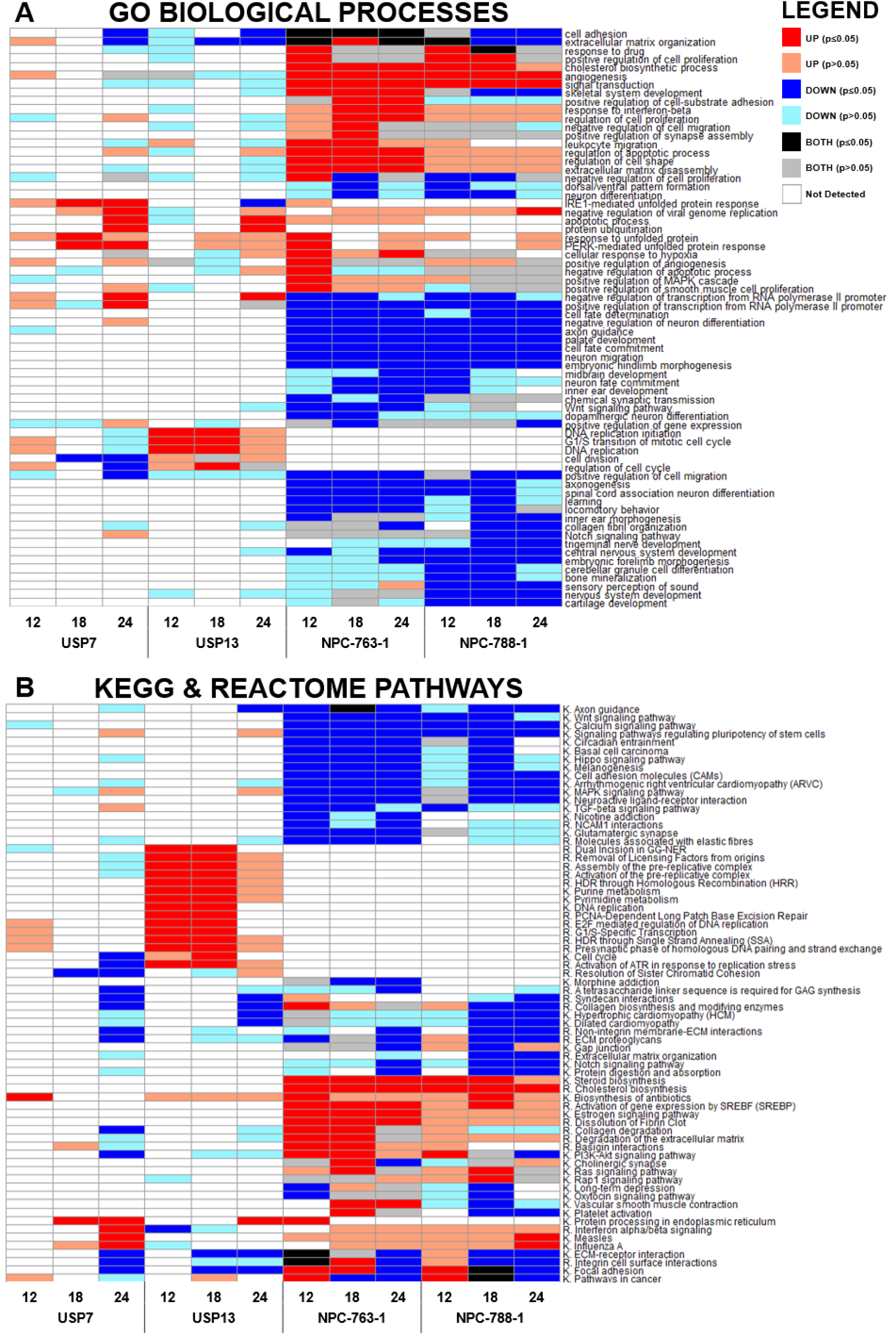
Differential biological processes and pathways following ZIKV infection. Heatmaps to show the **(A)** Gene Ontology (GO) Biological Processes and **(B)** KEGG and REACTOME Pathways that are differentially regulated following 12, 18 and 24 hours of ZIKV infection in brain tumour and neural precursor cells. KEGG and REACTOME pathways terms are labelled with the prefix of K. or R., respectively. All terms which were significant (p ≤ 0.05) in at least one condition were plotted, and non-significant (p > 0.05) enrichment for these terms across the other conditions was included. If a given term was enriched in both the upregulated and downregulated DEG lists, it was labelled BOTH. Significance values were corrected for multiple testing using the Benjamini and Hochberg method (padj ≤ 0.05). Abbreviations, Zika virus (ZIKV), USP7 (P7), USP13 (P13), NPC-763-1 (N3), NPC-788-1 (N7), Gene Ontology (GO), KEGG (K.), REACTOME (R.), hours post-infection (hpi), adjusted p-value (padj).

Comparing the largest DEGs at 24 hpi, we sought to identify the main conserved molecular mechanisms between the brain tumour cell lines. We observed a 34.2% overlap for brain tumour cell lines, a 42.5% overlap for NPCs, and no overlap between brain tumour cells and NPCs (Figure 3A). Plotting the 41 shared brain tumour DEGs (Figure 3B) and 51 shared NPC DEGs (Figure 3C) highlights a conserved cell-type specific transcriptome response to ZIKV infection as the directionality of all DEGs plotted are the same. KEGG pathway analysis on the top brain tumour DEGs identified ten significantly enriched terms that are predominantly involved in pathogen-host interaction (Viral protein interaction with cytokine and cytokine receptor, Toll-like receptor signalling pathway, Influenza A and Amoebiasis), immune response (Cytokine-cytokine receptor interaction, TNF signalling pathway, NF-kappa B signalling pathway and IL-17 signalling pathway) and cell death (Apoptosis) (Figure 3D). Considering the 41 shared brain tumour DEGs (Figure 3B), we sought to gain further insight into these enriched pathways to identify candidates for validation. We identify genes involved in TNF signalling and its downstream activity (TNF, TNFSF9, TNFRSF9, CCL5, MAP3K8, TRIB3, NFKBIE and OTUD1), the unfolded protein response (UPR) (ERN1, DDIT3, DNAJB9, ATF3 and HERPUD1), cell death (DDIT3, PMAIP1, BBC3, and CHAC1), antiviral responses (IFIT1, IFIT2, RSAD2, IFIH1 and OASL), interleukins (IL1A, IL7R and NFIL3), neuronal processes (RND1, LRRN3 and UNC5B), and lipid homeostasis (STARD4 and CH25H) (Figure 3B). Notably, we observe joint upregulation of the UPR-regulated pro-apoptotic transcription factor DDIT3 (CHOP), its downstream pro-apoptotic gene CHAC1 and BCL2-homologous (BH)-3-only proteins that are effectors of canonical mitochondrial apoptosis (PMAIP1 (NOXA) and BBC3 (PUMA)) (Figure 3B). These pro-apoptotic signatures and ER stress indicate ZIKV as a Type II ICD inducer for brain tumour cells, as expected for an ER-tropic virus. TNF and pro-inflammatory cytokine signalling are principal outcomes of ICD, which we observe as the top two ZIKV-infected brain tumour-enriched KEGG pathways (Figure 3D). To conclude, (i) ZIKV infection leads to conserved cell type-specific transcriptomic responses, (ii) NPCs are highly sensitive to ZIKV and undergo vast downregulation of neurodevelopmental processes following infection, and (iii) ZIKV-infected brain tumour cells portray ICD transcriptomic signatures.

**Figure 3.**
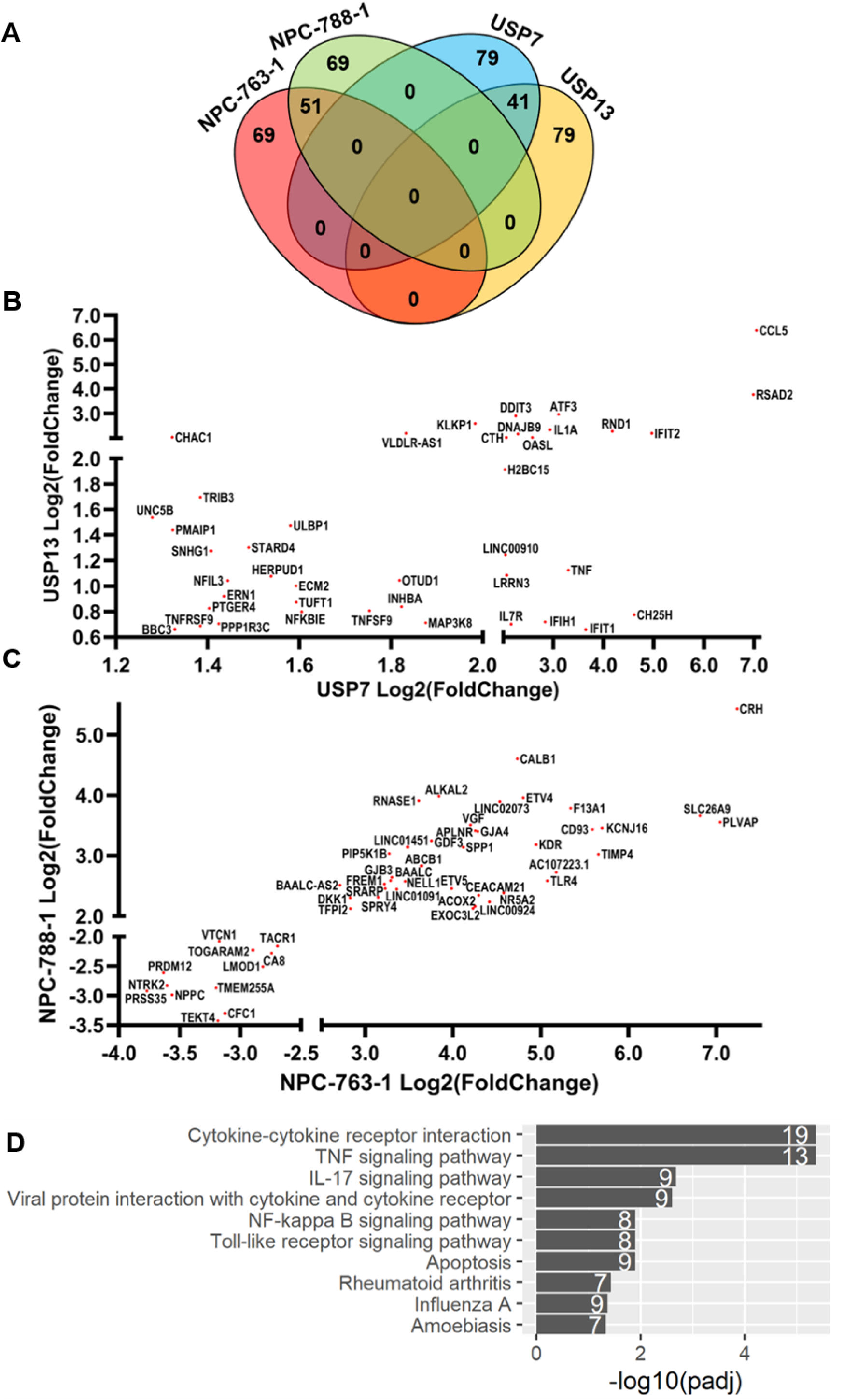
Cell type-specific responses in the top DEGs following ZIKV infection. **(A)** Venn diagram of the Top DEGs in each cell line at 24 hpi. Scatter plots of RNA-Seq Log2(Fold Change) for **(B)** the 41 overlapping brain tumour DEGs and **(C)** the 51 overlapping NPC DEGs at 24 hpi. **(D)** Barplot of KEGG pathways significantly enriched in the top brain tumour DEGs at 24 hpi. The white numbers at the end of each bar denote the number of DEGs identified for the given term. Significance values were corrected for multiple testing using the Benjamini and Hochberg method (padj ≤ 0.05). Abbreviations, adjusted p-value (padj).

### TNF-alpha is a potential prognostic marker for brain tumour ZIKV OV therapy

Assessing all differentially expressed TNF ligands, receptors, and adaptors, we observe the most highly upregulated DEGs in both ZIKV-infected USP7 and USP13 cells to be key components of the canonical TNF pathway (TNF-alpha) and non-canonical TNF receptor superfamily member 9 pathway (TNFSF9, TNFRSF9 and TRAF1) (Figure 4A). Activation of either pathway results in downstream NF-kappa B signalling, which we observe to be significantly enriched in the top ZIKV-infected brain tumour DEGs (Figure 3D). Data mining publicly available ZIKV-infected glioma and neuroblastoma RNA-Seq datasets, we observe that TNFRSF9 and one of its adaptors (TRAF1) are significantly upregulated in all cases, TNFSF9 is significantly upregulated in three instances, and TNF-alpha is significantly upregulated in all four brain tumour datasets (Figure 4B). Thus, these responses appear conserved across different ZIKV-infected nervous system tumour cells. Performing ProcartaPlex ELISA assays, we assessed whether the soluble TNF factors of these pathways (TNF-alpha, TNFSF9 and TNFRSF9) were secreted following infection and observed significantly increased secretion of TNF-alpha (Figure 4C) and TNFRSF9 (Figure 4D) from both USP7 and USP13 cells at 48 hpi. TNFSF9 was not reliably detected or quantified under any condition (Figure 4E). Thus, we show that ZIKV infection of brain tumour cells leads to the upregulated expression and secretion of key members of both canonical and non-canonical TNF signalling pathways.

**Figure 4.**
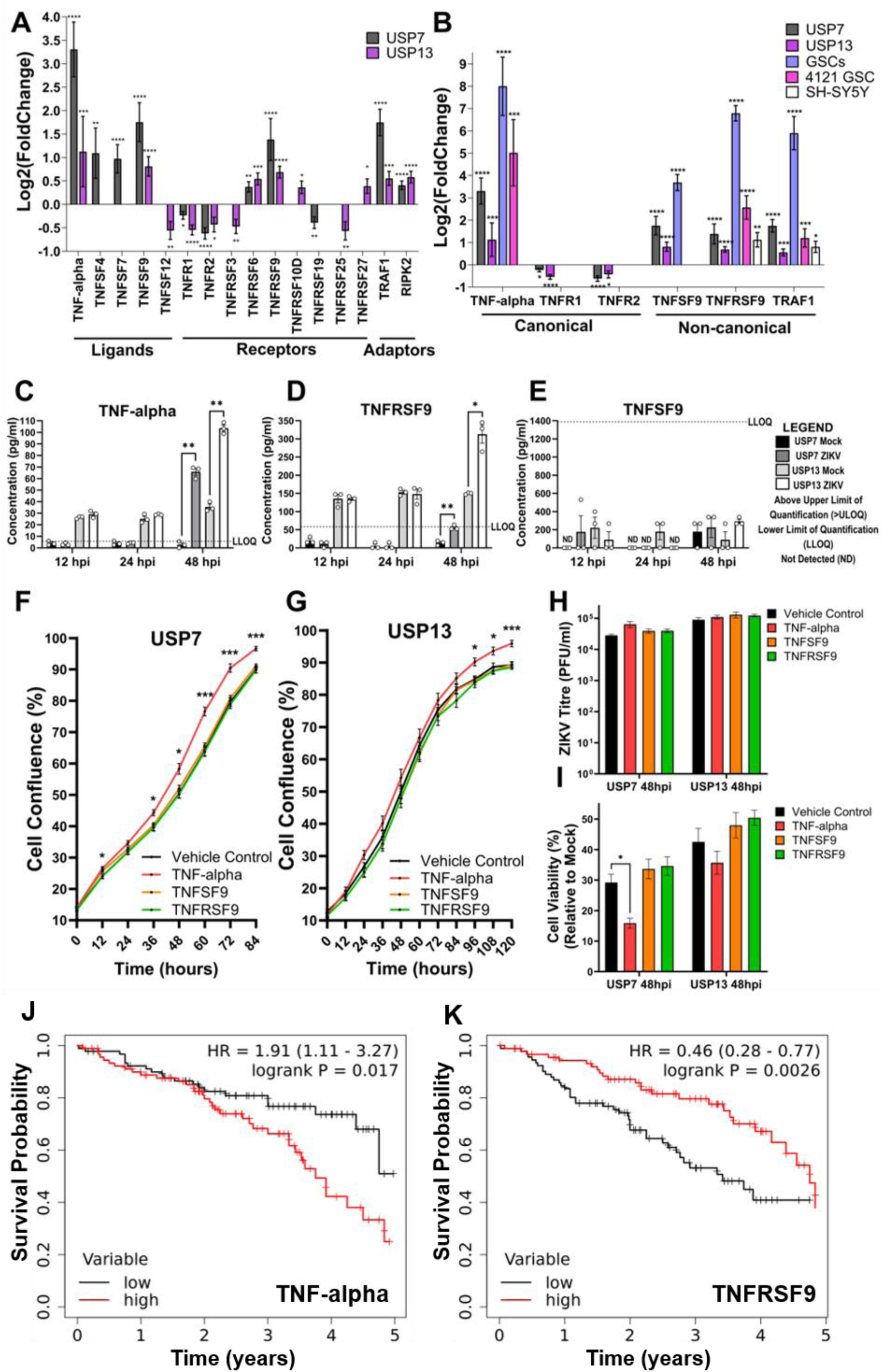
TNF signalling pathways during ZIKV infection of brain tumour cells. **(A)** Differentially expressed TNF pathway ligands, receptors, and receptor adaptors in ZIKV- infected USP7 and USP13 brain tumour cells at 24 hpi (LFC ± lfcSE, N=3). **(B)** Differential expression of principal genes of the canonical TNF-alpha and non-canonical TNFSF9/TNFRSF9 signalling pathways in nervous system tumour cells following ZIKV infection. ZIKV-infected RNA- Seq datasets include that generated in the current study for USP7 and USP13, and three sourced from GEO2R for GSCs, 4121 GSCs and SH-SY5Y neuroblastoma cells (LFC ± lfcSE, N=3). Protein concentration of secreted **(C)** TNF-alpha, **(D)** TNFRSF9 and **(E)** TNFSF9 from USP7 and USP13 cells following 12, 24 and 48 hours of ZIKV infection (Mean ± SEM, N=3). **(F)** USP7 and **(G)** USP13 cell confluence following recombinant TNF protein treatment (Mean ± SEM, n=8). **(H)** ZIKV titre following recombinant TNF protein treatment and ZIKV infection of USP7 and USP13 cells (Mean ± SEM, N=3, n=3). **(I)** USP7 and USP13 cell viability following recombinant TNF protein treatment and ZIKV infection (Mean ± SEM, N=3, n=3). Kaplan Meier plots to assess the correlation between medulloblastoma patient survival and the upper vs lower quartiles of **(J)** TNF-alpha and **(K)** TNFRSF9 gene expression. Logrank P and hazard rate (HR) with 95% confidence intervals are reported. Significance values were corrected for multiple testing using the Benjamini and Hochberg method for **A-B** (padj ≤ 0.05) and **C-E** (FDR ≤ 0.05) and Dunnett’s method for **F-I** (padj ≤ 0.05). Asterisk symbol denotes level of significance: *padj ≤ 0.05; **padj ≤ 0.01; ***padj ≤ 0.001; ****padj ≤ 0.0001. Abbreviations, Zika virus (ZIKV), Glioma Stem Cell (GSC), Log2(Fold Change) (LFC), standard error of the LFC estimate (lfcSE), Above Limit of Quantification (>ULOQ), Lower Limit of Quantification (LLOQ), Not Detected (ND), hours post-infection (hpi), adjusted p-value (padj).

Next, we sought to functionally assess these soluble proteins for their (i) effect on brain tumour cell growth, (ii) anti- or pro-viral properties, and (iii) effect on brain tumour cell viability following ZIKV infection. We employed Incucyte Live Cell Imaging to analyse USP7 and USP13 cell growth by tracking cell confluence following treatment with recombinant TNF proteins (Figure 4F and 4G). None of the recombinant TNF proteins were cytotoxic, and TNF-alpha significantly increased both USP7 and USP13 confluence, indicating that soluble TNF-alpha enhances brain tumour cell growth (Figure 4F and 4G). Next, we assessed the effect of recombinant protein treatment on ZIKV titre (Figure 4H) and on tumour cell viability (Figure 4I). Viral titre was unaffected by all TNF proteins, suggesting no anti- or pro-viral properties (Figure 4H). TNF-alpha significantly reduced ZIKV-infected USP7 cell viability and a modest but non-significant reduction was observed for ZIKV-infected USP13 cells (Figure 4I). Performing a five year survival analysis for 375 medulloblastoma patients clinical and gene expression data, we observe high TNF-alpha expression to be significantly correlated with worse survival (Figure 4J) and high TNFRSF9 expression to be significantly correlated with improved survival (Figure 4K). Thus, TNF-alpha is a marker of poor prognostic for medulloblastoma, possibly by enhancing brain tumour cell growth. Here, we show that TNF-alpha (i) is a brain tumour cell growth factor, (ii) may sensitise brain tumour cells to oncolysis by augmenting ZIKV-induced reduced cell viability, and (iii) is a marker of poor prognostic for medulloblastoma. Collectively, our results indicate that TNF-alpha holds promise as a potential prognostic marker, where a greater expression level of tumoural TNF-alpha may infer enhanced patient response to brain tumour ZIKV OV therapy.

### ZIKV infection stimulates a diverse pro-inflammatory cytokine secretome from brain tumour cells

Extending our TNF ELISA assays to a 49-plex ELISA, we sought to investigate the cytokine-related transcriptomic signatures enriched in ZIKV-infected brain tumour cells (Figure 3D). Assessing the temporal secretion of 34 cytokines (including TNF-alpha) and 15 immune checkpoint proteins (including TNFRSF9 and TNFSF9), we generate the most comprehensive ZIKV-infected cancer cell secretome to date (Supplementary Table 1). PCA clusters the secretome by cell line and highlights variation in the 48-hour ZIKV-infected condition for both cell lines from all other samples (Figure 5A). In addition to TNF-alpha (Figure 4C), we observe significantly increased secretion of 18 cytokines, predominately at 48 hpi (Figure 5B). Seven cytokines are significantly secreted by both infected cells (CXCL1, IL-1 alpha, IL-6, CXCL10 (IP- 10), CCL3, CCL4 and CCL5 (RANTES)), six from infected USP7 cells only (CCL11, IL-4, IL-8 (CXCL8), IL-9, IL-21 and SDF-1 alpha) and five from infected USP13 cells only (GM-CSF, IFN alpha, IL-1 beta, IL-1RA and IL-5) (Figure 5B). Conversely, of the 15 immune checkpoint proteins, TNFRSF9 was the only significantly secreted protein (Figure 4D), and PD-L2 was the only other protein secreted across all conditions (Supplementary Table 1). ZIKV infection did not significantly reduce the secretion of any protein. Integrating the 49-plex ELISA and RNA-Seq data, we observe 27 of the 49-plex proteins to be significantly differently expressed or secreted by USP7 or USP13 following ZIKV infection (Figure 6A). These diverse proteins encompass chemokine, interleukin, tumor necrosis factor, interferon, colony stimulating factor and programmed cell death proteins. Considering the primary role of each protein during an immune response, we identify that ZIKV infection predominantly promotes a pro-inflammatory rather than anti-inflammatory cytokine profile (Figure 6A). Additionally, we identify many cytokines which orchestrate adaptive immune responses. Crucially, many of these proteins are current OV transgenes or immunotherapy candidates. To conclude, ZIKV induces a diverse pro-inflammatory secretome from brain tumour cells, containing many clinically relevant candidates for mediating an OV-stimulated anti-tumoural immune response.

**Figure 5.**
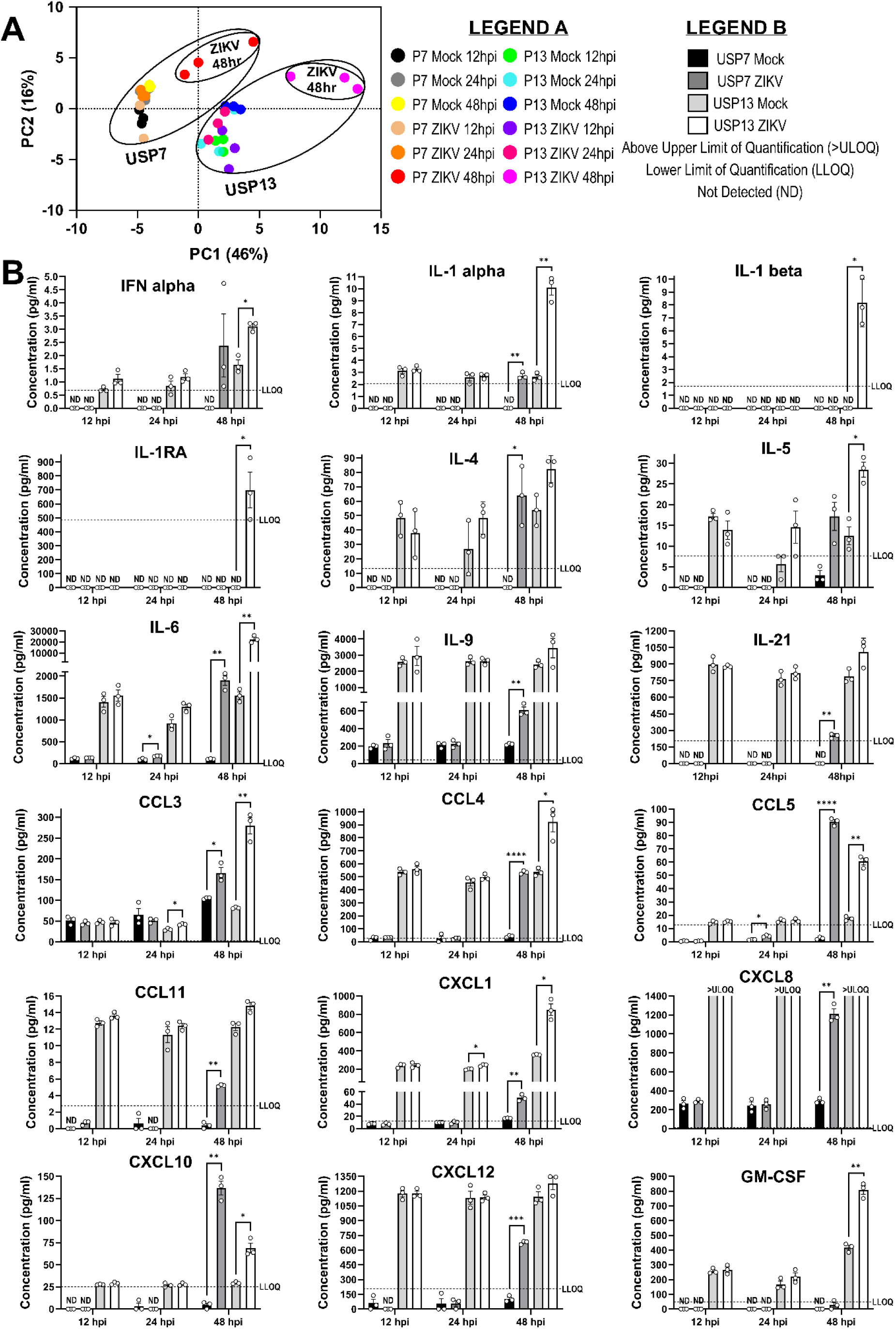
ZIKV-infected brain tumour cell secretome. **(A)** PCA plot of the 49-plex ELISA averaged Net MFI values (N=3). **(B)** Differential secretion of cytokines from USP7 and USP13 cells following 12, 24 and 48 hours of ZIKV infection (Mean ± SEM, N=3). Significance values were corrected for multiple testing using the Benjamini and Hochberg method (FDR ≤ 0.05). Asterisk symbol denotes level of significance: *padj ≤ 0.05; **padj ≤ 0.01; ***padj ≤ 0.001; ****padj ≤ 0.0001. Abbreviations, Zika virus (ZIKV), USP7 (P7), USP13 (P13), Above Limit of Quantification (>ULOQ), Lower Limit of Quantification (LLOQ), Not Detected (ND), hours post-infection (hpi), Principal Component (PC).

**Figure 6.**
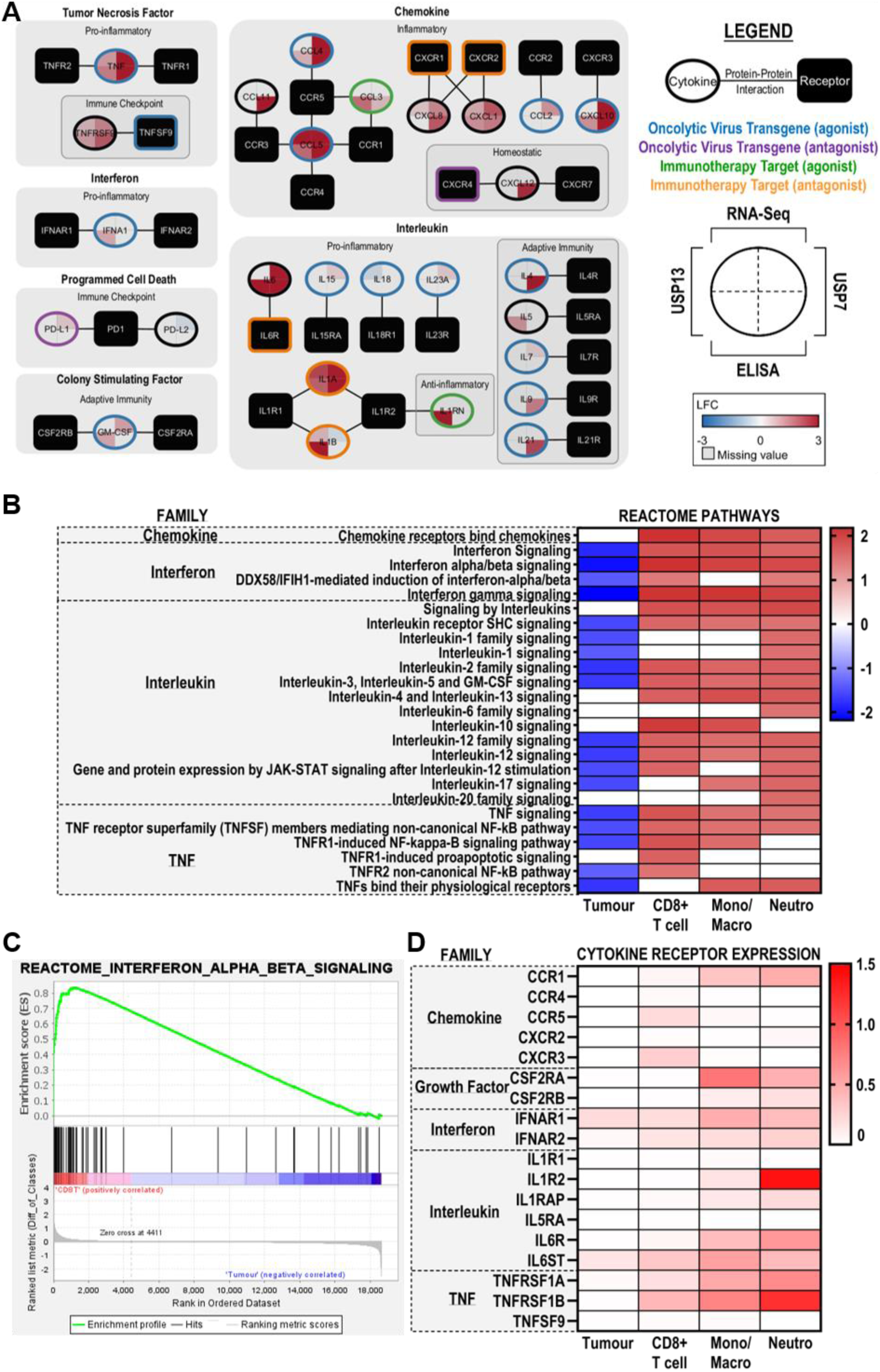
The pro-inflammatory ZIKV secretome and its predicted medulloblastoma TME paracrine signalling. **(A)** An integrated map of the 27 significantly expressed or secreted cytokines in at least one brain tumour cell line following ZIKV infection. Cytokines and their cognate receptors are clustered by their family class and sub-clustered by their primary role in the immune response. Edges denote cytokine-cytokine receptor interactions. Node shape denotes if the protein is a cytokine (oval) or receptor (rectangle). Piechart quadrants denote the maximum LFC value observed for the given cell line (USP7 or USP13) in the given assay (RNA-Seq or ELISA). Border colour denotes if the cytokine-receptor signalling is a current target of oncolytic virotherapy or immunotherapy and highlights which of the two proteins is the transgene or immunotherapy target. Heatmaps showing the **(B)** GSEA normalised enrichment score (NES) of significantly enriched cytokine-related REACTOME pathways (p ≤ 0.05, FDR ≤ 0.25) and **(D)** cytokine receptor expression (log-transformed normalised gene count) in the four major cell types annotated in the medulloblastoma scRNA-Seq. Rows for **(B)** and **(D)** are arranged by cytokine family. **(C)** Representative positively enriched cytokine-related GSEA enrichment plot (Interferon alpha/beta signalling) in immune cells (CD8+ T cells). Abbreviations, Zika virus (ZIKV), Log2(Fold Change) (LFC), Tumour microenvironment (TME), CD8+ T cells (CD8T), Monocyte/Macrophage (Mono/Macro), Neutrophil (Neutro), Gene Set Enrichment Analysis (GSEA), Normalised Enrichment Score (NES).

### The medulloblastoma tumour immune microenvironment (TIME) is primed to respond to pro-inflammatory ZIKV secretome paracrine signalling

Employing an *in silico* approach, we sought to model how the ZIKV-induced pro-inflammatory USP13 medulloblastoma secretome may interact with tumour and immune cells of the medulloblastoma TME. Analysing publicly available medulloblastoma scRNA-Seq, we identify positive enrichment of many cytokine-related REACTOME pathways by GSEA (Figure 6B and 6C) and greater cytokine receptor expression (Figure 6D) in monocytes/macrophages, CD8+ T cells and neutrophils compared to medulloblastoma tumour cells. All immune cells express the TNF-alpha (TNFRSF1A and TNFRSF1B) and IFN alpha (IFNAR1 and IFNAR2) receptors and are enriched for these pathways, indicating they are primed to respond to secreted TNF-alpha and IFN alpha in the TME (Figure 6B-D). Monocytes/macrophages and neutrophils express the CCL3 and CCL5 receptor CCR1, while CD8+ T cells express the CXCL10 receptor CXCR3 and the CCL4 and CCL5 receptor CCR5. All three immune cell types are enriched for the Chemokine receptors bind chemokines REACTOME pathway, and thus likely respond to these secreted chemokines. Additionally, monocytes/macrophages and neutrophils express the main GM-CSF receptor CSF2RA and are enriched in the IL-3, IL-5 and GM-CSF signalling pathway, indicating potential to respond to secreted GM-CSF. Finally, neutrophils express the IL-1 (IL1R2 and IL1RAP) and IL-6 (IL6R and IL6ST) receptors and are the only cell type enriched for these pathways, indicating a possible neutrophil specific response to ZIKV-induced medulloblastoma-secreted interleukins. Notably, medulloblastoma tumour cells are not positively enriched for any REACTOME cytokine-related pathway and only express two receptors (IFNAR1 and IL6ST) at low levels, indicating that they are less likely to respond to the ZIKV-induced medulloblastoma secretome (Figure 6B and 6D). Collectively, these observations suggest that the ZIKV-induced pro-inflammatory secretome is more likely to target TME immune cells to remodel the TIME rather than non-infected medulloblastoma tumour cells.

### ZIKV secretome endocrine signalling is predicted to stimulate diverse immune cell polarisations

Integrating our ZIKV-induced brain tumour secretome with the Immune Dictionary, we next modelled how endocrine signalling may induce immune cell polarisations towards anti-tumoural activation states following oncolytic ZIKV infection. Six of our 20 differentially secreted proteins (IFN alpha, IL-1 alpha, IL-1 beta, IL-4, TNF-alpha and GM-CSF) are documented in the Immune Dictionary as dominant cytokines which drive polarisation states of the main lymphoid (CD4+ T, CD8+ T and NK cells) and myeloid (monocytes, macrophages, dendritic cells (DCs) and neutrophils) cell types implicated in anti-tumoural immune responses (36). Assessing individual cytokine-immune cell interactions, the resulting polarisation states, and the top-upregulated DEGs, we sought to elucidate the immune cell phenotypes following cytokine stimulation (Figure 7). We observe all six cytokines to be pleiotropic, polarising the eight immune cell types into 26 distinct states (Figure 7). IFN alpha, IL-1 alpha, and IL-1 beta polarise every cell type. IFN alpha induces a unique polarisation (suffix -a), and IL-1 alpha and IL-1 beta (collectively referred to as IL-1) induce a conserved polarisation (suffix -c). We observe self-regulation of cytokine signalling transduction by positive feedback receptor upregulation (e.g. IL4R), negative feedback inhibitory receptor upregulation (e.g. IL1R2), and upregulation of cytokine signalling regulators (SOCS1, SOCS3, CISH and DUSP10). Most cytokines sensitise or suppress the immune cells to additional cytokine signalling through upregulated expression of cytokines (e.g. CXCL9), receptors (e.g. IFNGR1), or receptor antagonists (e.g. IL1RN).

**Figure 7.**
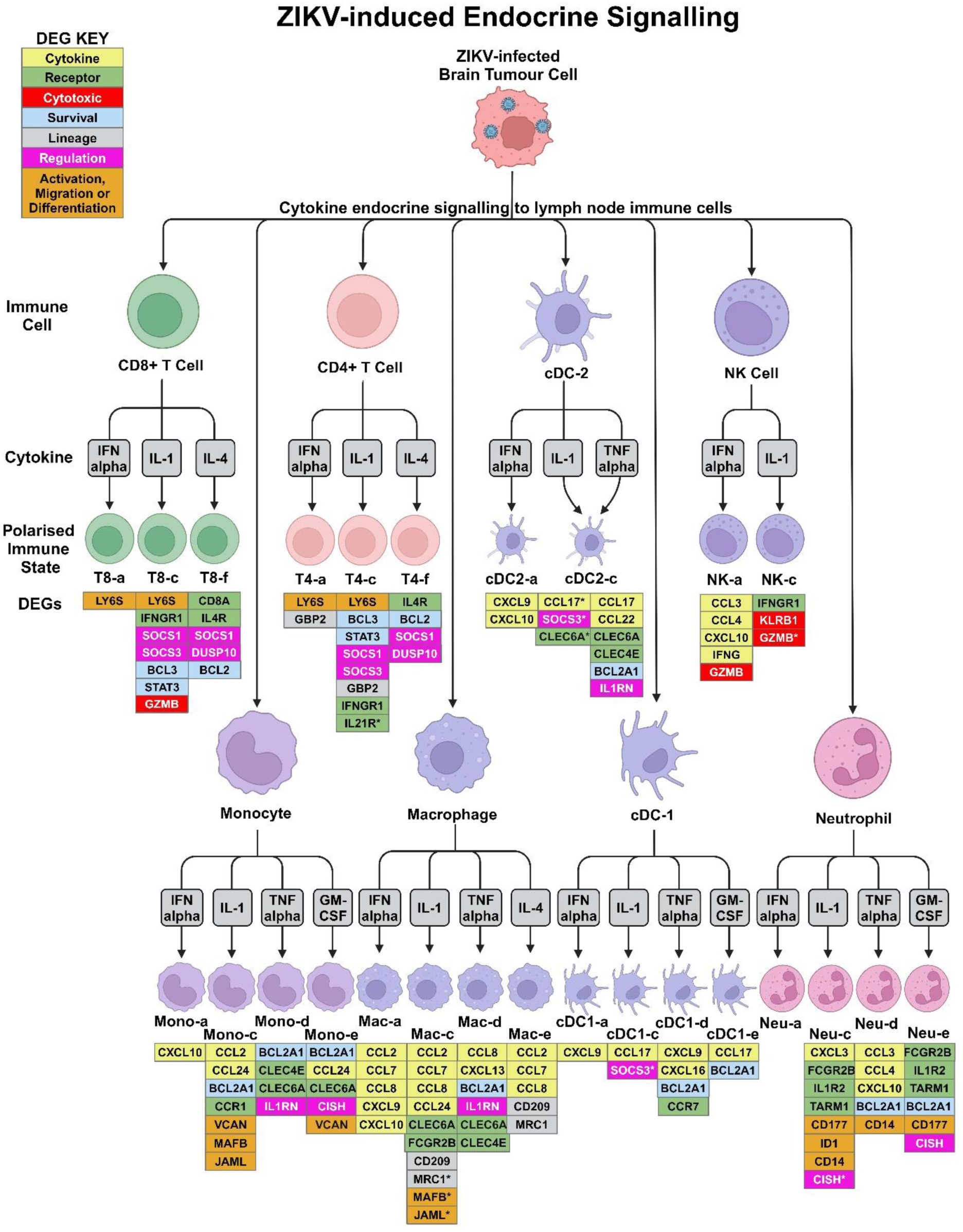
Modelling of ZIKV-induced secretome endocrine signalling to lymph node immune cells. Tree diagram detailing polarisation states of lymph node immune cells following cytokine stimulation, as found in the Immune Dictionary (36). Only ZIKV-induced brain tumour cell-secreted cytokines that are dominant drivers of immune cell polarisation were assessed, as determined by the Immune Dictionary. Polarisation states, their nomenclature and the top 20 upregulated cytokine-induced DEGs (FDR ≤ 0.05) were sourced from the Immune Dictionary. Phenotypic DEG markers that indicate immune cell activity or phenotype following cytokine stimulation are listed below the respective polarised state, with colours denoting if the DEG is a cytokine (yellow), receptor (green), cytotoxicity-related gene (red), survival-related gene (blue), cell lineage marker (grey), cytokine signalling regulator (pink), or a gene indicating immune cell activity (activation, migration or differentiation) (orange). IL-1 alpha and IL-1 beta induce the same polarisation state and are thus collectively represented as IL-1. * denotes a DEG induced by either IL-1 alpha or IL-1 beta, but not both. Abbreviations, Zika virus (ZIKV), conventional Dendritic Cell (cDC), Natural Killer (NK), Differentially Expressed Gene (DEG), CD8+ T cell (T8), CD4+ T cell (T4), Monocyte (Mono), Macrophage (Macro), Neutrophil (Neu). Figure created in BioRender. BioRender.com/u63c617.

We observe TNF-alpha to act on myeloid rather than lymphoid lineage cells (Figure 7). All TNF-alpha and GM-CSF-stimulated myeloid cells upregulate the anti-apoptotic BCL2A1 gene, indicating a BCL2A1-mediated pro-survival mechanism. TNF-alpha induces the expression of IL1RN (IL-1RA) and the pattern recognition receptors CLEC6A and CLEC4E in monocytes, macrophages and cDC2s, possibly suppressing IL-1 responses and priming myeloid cells for innate immune recognition and activation. Upregulation of LY6S indicates CD4+ and CD8+ T cell activation by IFN alpha and IL-1 (T4-a, T4-c, T8-a and T8-c). Upregulation of the effector memory T cell marker GBP2 indicates that T4-a and T4-c likely become effector memory CD4+ T cells. Upregulation of the principal cytotoxic granzyme GZMB indicates that T8-c are likely cytotoxic CD8+ T cells. T cell survival may be driven in a BCL2-mediated manner following IL-4 stimulation (T4-f and T8-f) and a STAT3/BCL3-mediated manner following IL-1 stimulation (T4-c and T8-c). IFN alpha and IL-1 beta upregulate cytotoxic granzyme GZMB expression in NK-a and NK-c cells, and the cytotoxicity regulator KLRB1 may govern the NK-c cytotoxic state. As IFN gamma (IFNG) signalling promotes cytotoxicity, all observed cytotoxic polarisation will likely be augmented by their upregulation of IFNG (NK-a) or its receptor IFNGR1 (NK-c and T8-c). Notable upregulation of C-C Motif Chemokine Ligands (CCL) known to govern immune cell chemotaxis are observed for all macrophage polarisations, with a conserved CCR2 axis (CCL2, CCL7 and CCL8) for Mac-a, Mac-c and Mac-e. The Mac-c and Mac-e polarisations present with upregulated M2 macrophage marker (MRC1 (CD206) or CD209) expression, implicating them as M2 macrophages. Conversely, Mac-a presents with upregulated expression of pro-inflammatory CXCL9 and CXCL10 cytokines, indicative of M1 macrophages. IL-1 and GM-CSF-induced Mono-c and Mono-e polarisations upregulate M1 macrophage differentiation marker VCAN, suggesting monocyte to M1 macrophage differentiation. Both IL-1-induced Mono-c and IL-1 beta-induced Mac-c polarisations have upregulated monocyte/macrophage migration (JAML) and differentiation (MAFB) marker expression. To conclude, *in silico* modelling of ZIKV-induced secretome endocrine signalling indicates a diverse immune cell response, including (i) effector memory CD4+ T cells (T4-a and T4-c), (ii) cytotoxic CD8+ T cells (T8-c), (iii) cytotoxic NK cells (NK-a and NK-c), (iv) M1 (Mac-a) and M2 (Mac-c and Mac-e) macrophages, (v) M1 macrophage-destined monocytes (Mono-c and Mono-e), and (vi) neutrophils, cDC and other immune cell polarisations of currently unknown phenotype.

## Discussion

In the current study, we perform the first multi-omics-based investigation of ZIKV infection in paediatric brain tumour cells. At DEG, biological process and pathway levels, we demonstrate a stark contrast in the response of ZIKV-infected brain tumour cells to that of CZS patient-derived NPCs. ZIKV infection leads to a potent anti-tumoural immune response against mouse glioma, resulting in tumour clearance and long-term immunity against tumour cells (30–32). It is currently unknown what ICD mechanisms orchestrate this crucial efficacious response. Here, our multi-omics analysis highlighted the involvement of TNF and cytokine signalling as indicators of ICD in ZIKV-infected childhood brain tumour cells (37). Previous observations by our group support these observations and have shown that (i) this transcriptomic TNF and cytokine response is also observed in ZIKV-infected neuroblastoma cells, (ii) ZIKV-infected brain organoids co-cultured with USP7 or USP13 cells upregulate expression of TNF and a limited number of related cytokines, and (iii) ZIKV-infected canine glioblastoma cells secrete elevated levels of interleukins (28,29,33).

TNF signalling is a diverse signalling pathway capable of regulating cell proliferation, cell death and immune responses (38). Contrasting studies report TNF-alpha as anti-tumoural (cytotoxic and cytostatic) or pro-tumoural for medulloblastoma, and that both TNF-alpha transgene and TNFR antagonist treatment portray favourable outcomes in xenograft models (39–41). Regarding OVs, two independent groups working on the oncolytic myxoma virus report contrasting results that either TNF-alpha transgene or blockade enhances OV efficacy (42,43). Thus, the clinical utility of TNF-alpha in paediatric brain tumour therapy and OV therapy requires further delineation. Here, we observe TNF-alpha as not cytotoxic but pro-tumoural to medulloblastoma and ATRT cells under normal conditions. Additionally, we identify TNF-alpha as a marker of poor prognosis in medulloblastoma, likely in part due to enhanced tumour growth (38). Interestingly, recombinant TNF-alpha reduces cell viability of ZIKV-infected brain tumour cells, suggesting that in the context of ZIKV OV therapy, TNF-alpha is anti-tumoural as it enhances brain tumour cell oncolysis. This response is TNFR1-dependent, as TNFR2 activation requires membrane-bound TNF-alpha (44). Whilst we did not observe TNF-alpha to be anti- or pro-viral in medulloblastoma or ATRT cells, it is anti-viral in ZIKV-infected adult glioblastoma cells (45). Our *in silico* analysis indicates that all medulloblastoma TME immune cells are primed to respond to TNF-alpha paracrine signalling and that TNF-alpha endocrine signalling would polarise myeloid cells to upregulate pro-survival and innate immune recognition genes. Our observed anti-tumoural role of TNF-alpha during ZIKV oncolysis, its reported anti-viral property, and our *in silico* prediction of its involvement in anti-tumoural immune responses highlight that oncolytic ZIKV infection could likely be augmented by adjuvant therapy targeting TNF-alpha. This concept is exemplified by oncolytic HSV-1 inducing TNF-alpha-mediated glioma cell death and adjuvant TNF-alpha blockade enhancing both viral replication and mouse survival (46).

Pro-inflammatory cytokine secretion is a hallmark signature of ICD, and here, we demonstrate by multiplex ELISA that ZIKV induces a diverse pro-inflammatory secretome from childhood brain tumour cells. This natural induction of a diverse pro-inflammatory secretome is essential to ZIKV OV therapy efficacy because ZIKV’s limited RNA genome size is a barrier to enhancing this response by stable transgene modification. Currently, only a limited number of multiplex ELISA assays have assessed the secretome of ZIKV-infected brain tumour cells (26,33). A 25- plex ELISA, measuring 23 of the 34 cytokines we consider here, showed that ZIKV infection only significantly increased CXCL10 and CCL5 secretion and decreased CCL2 secretion from adult glioblastoma cells (26). The size and diversity of this cytokine response is much smaller than that we show for ZIKV-infected paediatric medulloblastoma and ATRT cells, suggesting that oncolytic ZIKV infection is more immunogenic for paediatric rather than adult brain tumour cells.

Here, we observe ZIKV infection to elevate secretion of the principal triad of pro-inflammatory cytokines (IL-1, IL-6, and TNF-alpha) by both brain tumour cell lines, in addition to many other cytokines that we report for the first time to be a component of oncolytic ZIKV infection. Corroborating our findings, three key pro-inflammatory cytokines with known anti-tumoural functions (IL-6, CCL5 and CXCL10) are significantly secreted by ZIKV-infected glioblastoma cells as well as ZIKV-infected USP7 and USP13 cells (26,33). We observe ZIKV to upregulate secretion of numerous drivers of OV-induced anti-tumoural immune responses, including but not limited to IFN alpha, IL-1, IL-6, CCL5, CXCL8, CXCL10, CXCL12, GM-CSF and TNF-alpha; as reviewed in (47). In addition to detecting the secretion of numerous cytokines that are OV transgenes or immunotherapy target candidates (Figure 6A), we observe ZIKV-infected USP13 cells to specifically upregulate IFN alpha and GM-CSF secretion, both of which are FDA-approved cytokine therapies (48). IFN alpha is a monotherapy for treating various adult cancers and GM-CSF is an adjuvant to high-risk neuroblastoma anti-GD2 immunotherapy (48–50). Here, we report that ZIKV induces a clinically diverse pro-inflammatory and anti-tumoural brain tumour secretome primarily composed of cytokines currently employed as OV transgenes, immunotherapy targets or cytokine therapies.

A crucial element of ZIKV OV efficacy is the ability to heat up the immunosuppressive TIME by paracrine signalling to co-opt TME immune cell activity and endocrine signalling to orchestrate immune cell infiltration. ZIKV-induced brain tumour immune cell infiltration includes CD4+ T cells, CD8+ T cells, NK cells, monocytes, macrophages, DCs and microglia cells (30–32). Modelling these *in silico*, we predict distinct polarisation states of the majority of these infiltrating immune cells in response to oncolytic ZIKV infection. Both CD8+ T cells and NK cells infiltrate ZIKV-infected mouse glioma, and here, our modelling indicates anti-tumoural cytotoxic CD8+ T and NK cell polarisations in response to ZIKV-induced brain tumour cell secretome endocrine signalling (30). This anti-tumoural phenotype is characterised by upregulated expression of the pro-apoptotic granzyme B, which is a principal component of cytotoxic granules, in addition to key markers of IFN gamma signalling, which orchestrate interferon-mediated anti-tumoural immune responses (51). ZIKV-induced tumour clearance is dependent on cytotoxic CD8+ T cells, and our modelling indicates that this cytotoxic polarisation is in response to IL-1 (30).

Tumour-associated macrophages (TAMs) generally exist in the pro-tumoural and immunosuppressive M2 state and commonly undergo a phenotypic shift to a pro-inflammatory and anti-tumoural M1 state following OV infection (52). M1 macrophage infiltration has been reported following ZIKV infection of mouse and canine brain tumours *in vivo*, however, the mechanisms that orchestrate this response are unknown (30,33). Of the 13 significantly secreted cytokines from ZIKV-infected USP13 medulloblastoma cells, nine (TNF-alpha, GM-CSF, IL-1 beta, IL-6, CXCL1, CXCL10 and CCL3-5) are known to induce the M1 phenotype, whilst only CCL5 and high levels of IL-1RA may lead to the M2 phenotype (53). Here, we observe high TNF-alpha and GM-CSF receptor expression and significant enrichment of their respective pathways in medulloblastoma TAMs. As such, we propose that in response to ZIKV-induced secretion of either TNF-alpha or GM-CSF, these medulloblastoma TAMs would adopt a pro-inflammatory and anti-tumoural phenotype. Utilising the Immune Dictionary, we identify that macrophages are stimulated by IFN alpha and monocytes are stimulated by both IL-1 and GM-CSF to express M1 macrophage gene markers. In contrast, IL-1 and IL-4 stimulate M2 gene markers in macrophages, which supports the known immunosuppressive effects of these cytokines on TAMs (54). Therefore, our modelling indicates a predominantly M1 macrophage phenotype following oncolytic ZIKV infection with potential for some M2 macrophage populations.

Coupling the ZIKV-induced pro-inflammatory cytokine profile with immunotherapy presents a unique opportunity to enhance efficacy by augmenting tumour destruction. ZIKV remodelling of the immunosuppressive TIME sensitises mouse glioma to PD-1 and PD-L1 immune checkpoint blockade (30,31). CXCL8 blockade sensitises adult glioma to anti-PD-1, and here we identify CXCL8 to be highly secreted by USP13 under all conditions and by ZIKV-infected USP7 cells at 48hpi (55). This may indicate a potential multi-modal therapy strategy, and multi-modal therapy to augment oncolytic ZIKV infection of paediatric brain tumours deeply warrants investigation.

To conclude, we investigate ICD signatures following oncolytic ZIKV infection of paediatric brain tumours and propose TNF-alpha as a potential prognostic marker for brain tumour ZIKV OV therapy, with the only other known prognostic marker being CD24 for neuroblastoma ZIKV OV therapy (56). We observe a clinically relevant pro-inflammatory and anti-tumoural ZIKV-induced brain tumour secretome. Through *in silico* modelling, we indicate that ZIKV secretome paracrine signalling likely targets TME immune cells over non-infected medulloblastoma tumour cells and endocrine signalling to drive diverse immune cell polarisations. We propose that this would collectively lead to a pro-inflammatory and anti-tumoural immune response that would heat up the paediatric brain TME and be fundamental to the efficacy of ZIKV OV therapy. The research we perform here contributes to understanding the molecular mechanisms governing oncolytic ZIKV infection and the cytokine response which orchestrates the anti-tumoural immune response. This work contributes to the growing body of research indicating the clinical utility of ZIKV as an oncolytic virotherapy for nervous system tumours and will help pave the way towards its application in clinical trials.

## Methods

### Cell and Viral Culture

Paediatric USP7 ATRT and USP13 medulloblastoma cells were cultured as previously described (21,57), as were the paediatric CZS-affected patient-derived NPCs (NPC-763-1 and NPC-788-1) (35). Vero cells were cultured in complete DMEM (Gibco, 41966029) with 10% FBS and 1% Penicillin-Streptomycin (5% CO_2_, 37°C) and passaged 1:12 every 72 hours. Stocks of two highly conserved Brazilian epidemic ZIKV strains, BeH819966 (KU365779.1) and Paraiba_01 (KX280026.1), were maintained at Instituto Butantan (University of São Paulo) and The Pirbright Institute, respectively. To maintain stocks, Vero cells in serum-free DMEM were infected for 90 min. At 96 hpi or when sufficient CPE was detected, virus was collected, clarified, filtered (0.22 μm) and stored as single use aliquots at −80°C. Virus was titrated in technical duplicate in Vero cells by plaque forming units (PFU) assay, using a 10-fold dilution series in serum-free DMEM. Following 90 min infection, cells were washed with PBS, a solid overlay of UltraPure 1% LMP agarose (Invitrogen, 16520-100) containing 0.83% complete DMEM was added, and plates were incubated for 72 hours. Plaques were fixed with formalin overnight, washed with PBS, and stained with crystal violet for 5 min. All brain tumour cell and NPC infection experiments were performed at a multiplicity of infection (MOI) of 2 for 60 min. Infection experiments for confocal microscopy and RNA-Seq used BeH819966, whilst all remaining infection experiments used Paraiba_01.

### Confocal Microscopy

Brain tumour cells were plated on coverslips for 24 hours prior to infection, infected and then fixed with 4% PFA for 40 min. Cells were permeabilised and blocked with PBSAT for 30 min, incubated in primary antibody anti-NS2B (Genetex, GTX133308) for 90 min, and then secondary antibody (ThermoFisher, A-11007) for 60 min. Cells mounted in VECTASHIELD Medium with DAPI (Vector Laboratories, H-1200-10). Coverslips were washed with PBS three times between incubations. Coverslips were stored until imaged via confocal microscopy with ZEN Software.

### RNA-Sequencing

At 12, 18 and 24 hpi, cells were collected, washed with PBS and stored as pellets at −80°C (N=3). High-quality total RNA was purified using the Monarch Total RNA Miniprep Kit (NEB #T2010S) per the manufacturer’s protocol. mRNA sequencing was performed by Novogene (UK) Company Limited using the Illumina NovaSeq 6000 system (≥20 million 150 bp paired-end reads per sample). The RNA-Seq data processing pipeline consisted of FastQC (V0.11.9-0), Trim Galore (V0.6.6-0), HISAT2 (V2.2.0), Samtools (V1.11) and Subread (V2.0.1). Reads were aligned against the Homo Sapien GRCh38 and ZIKV BeH819966 genome, and differential gene expression analysis was performed using DESeq2 to identify brain tumour (padj ≤ 0.05, Fold Change ≥ 0) and NPC (padj ≤ 0.05, Fold Change ≥ 1.5) DEGs. Upregulated and downregulated DEGs were submitted to DAVID to identify upregulated and downregulated GO Biological Processes, KEGG pathways and REACTOME pathways, respectively (58). Significance values were corrected for multiple testing using the Benjamini and Hochberg method (padj ≤ 0.05). Hierarchical clustering ordered the terms (rows) by similarity across the twelve infection conditions. The top 120 DEGs were determined by ranked absolute Log2(Fold Change) (LFC) value, and KEGG pathway analysis was performed for these brain tumour DEGs using DAVID. The RNA-Seq data generated in the present study have been deposited to the NCBI Gene Expression Omnibus (GEO) with the GEO accession GSE277900. The publicly available RNA-Seq datasets of ZIKV-infected tumour cells were sourced from GEO2R with the GEO accessions GSE114907 (4121 GSC, MOI 1, 48 hpi, N=3), GSE102924 (387, 3565 and 4121 GSC, MOI 5, 48 hpi, N=3) and GSE149775 (SH-SY5Y, MOI 5, 24 hpi, N=3) (24,26,59). All DEGs are presented as LFC with p-values corrected for multiple testing by the Benjamini and Hochberg method (padj ≤ 0.05).

### Recombinant TNF Assays

USP7 and USP13 cells were treated with 100µg/ml recombinant TNF-alpha (Peprotech, 300- 01A), TNFSF9 (Peprotech, 310-11), TNFRSF9 (Peprotech, 310-15) or a 0.1% BSA vehicle control. For Incucyte Live Cell Growth Analysis, USP7 and USP13 cells were plated at low confluence in sufficient media and allowed to grow until the first condition of each cell line reached 95% confluency. Confluence was determined by the Confluence Basic Analyser, with the phase map trained using AI specifically off each cell line (label-free) at varying confluency to ensure the accuracy of readings during the live cell imaging. Cell culture supernatant was collected at 48 hpi for infection experiments, clarified, stored as single-use aliquots at −80°C, and titrated via PFU in Vero cells. Cell viability was determined using the CellTiter-Glo Assay (Promega, G7572), per the manufacturer’s protocol. For all recombinant TNF assays, significance was defined and corrected for multiple testing by two-way ANOVA with Dunnett’s multiple correction (padj ≤ 0.05).

### Medulloblastoma Patient Survival Analysis

Medulloblastoma patient gene expression and survival data were sourced using GEO2R for GEO accession GSE85218 (GPL22286) (60). ATRT patient survival analysis could not be performed due to the extreme scarcity of ATRT datasets. Patient survival was censored at five years, and Kaplan-Meier survival analysis was performed for the remaining 375 patients. Logrank P was determined by univariate cox regression (p ≤ 0.05), comparing survival between the upper and lower quartiles based on candidate gene expression. Hazard rate (HR) with 95% confidence intervals is reported. Analysis was performed using KMplot (61).

### ProcartaPlex Multiplex ELISA Assay

At 12, 24 and 48 hpi, cell culture supernatant was collected from infected USP7 and USP13 cells, centrifuged, and stored as single-use aliquots at −80°C (n=2, N=3). ProcartaPlex kits quantified supernatant protein concentration of 34 cytokines (Invitrogen, EPXR340-12167-901) and 15 immune checkpoint proteins (Invitrogen, EPX010-15901-901 and EPX14A-15803-901), as per the manufacturer’s protocol. Data was collected using a Bio-Plex 200 and processed on the ThermoFisher ProcartaPlex Analysis App. Optimised standard curves were plotted with a 5 PL Logistic fit. For each condition, outliers were omitted and technical duplicates were averaged. Multiple unpaired t-tests defined significance with Benjamini and Hochberg correction (FDR ≤ 0.05). PCA was performed on the averaged Net MFI values.

### Paracrine Signalling Analysis

scRNA-Seq log-transformed normalised gene counts matrix of 28 untreated primary paediatric human medulloblastoma tumours were sourced from the Tumor Immune Single-cell Hub 2 (TISCH2) (62). Analysis was limited to medulloblastoma due to the absence of ATRT scRNA-Seq datasets. The scRNA-Seq dataset consisted of 37,445 cells: 32,926 Tumour cells, 2,665 monocytes/macrophages, 1,590 CD8+ T cells and 264 neutrophils (GEO accession GSE155446) (63). TISCH2 processed and analysed the dataset using the MAESTRO workflow, and the gene counts matrix reports the average expression per cell type. GSEA and the expression of specific cytokine receptors of interest were plotted using the log-transformed normalised gene counts matrix. GSEA was performed by comparing each immune cell type against the tumour cells for immune cell pathway enrichment, and then tumour cells against all immune cells for tumour cell pathway enrichment (64). Analysis was performed with the REACTOME database, and significant pathways (p ≤ 0.05 and FDR ≤ 0.25) relating to Interleukin-, Interferon-, TNF- and Chemokine-receptor signalling were plotted.

### Endocrine Signalling Analysis

Dominant cytokine drivers of immune cell polarisation were identified from the Immune Dictionary publication (36). scRNA-Seq of mouse lymph node cells following individual recombinant cytokine treatment *in vivo* was sourced using the Immune Dictionary Application (IDA). Within the IDA, DEGs were determined by a two-sided Wilcoxon rank-sum test comparing normalised gene expression values of cytokine treatment versus PBS control (N=3), with significance corrected for multiple testing (FDR ≤ 0.05). The Top 20 upregulated DEGs for each cytokine-immune cell interaction of interest (Supplementary Table 2) were used to infer the cytokine-induced immune cell polarisation phenotype. DEGs of interest were mapped to their human orthologs using g:Profiler for inclusion in Figure 7 (65).

## Acknowledgements

The authors acknowledge financial support from UKRI (MR/S01411X/1) (to RME and OKO), Wessex Medical Research Trust (to RME), Rosetrees Trust (to RME), The Little Princess Trust (to RME), FAPESP-CEPID (Proc. No. 2013/08028-1) (to OKO), and CNPq (Proc. No. 302736/2022-0)(to OKO) for this project. KM receives salary and research support from BBSRC grants BBS/E/PI/230001A and BBS/E/PI/23NB0003. The authors would like to thank the Elma Tchilian research group at The Pirbright Institute for sharing their Bio-Plex 200 calibration and validation kits, and Ehsan Rostami in particular for his support in setting up the 49-plex ProcartaPlex ELISA assay.

---

**Supplementary Figure 1.**
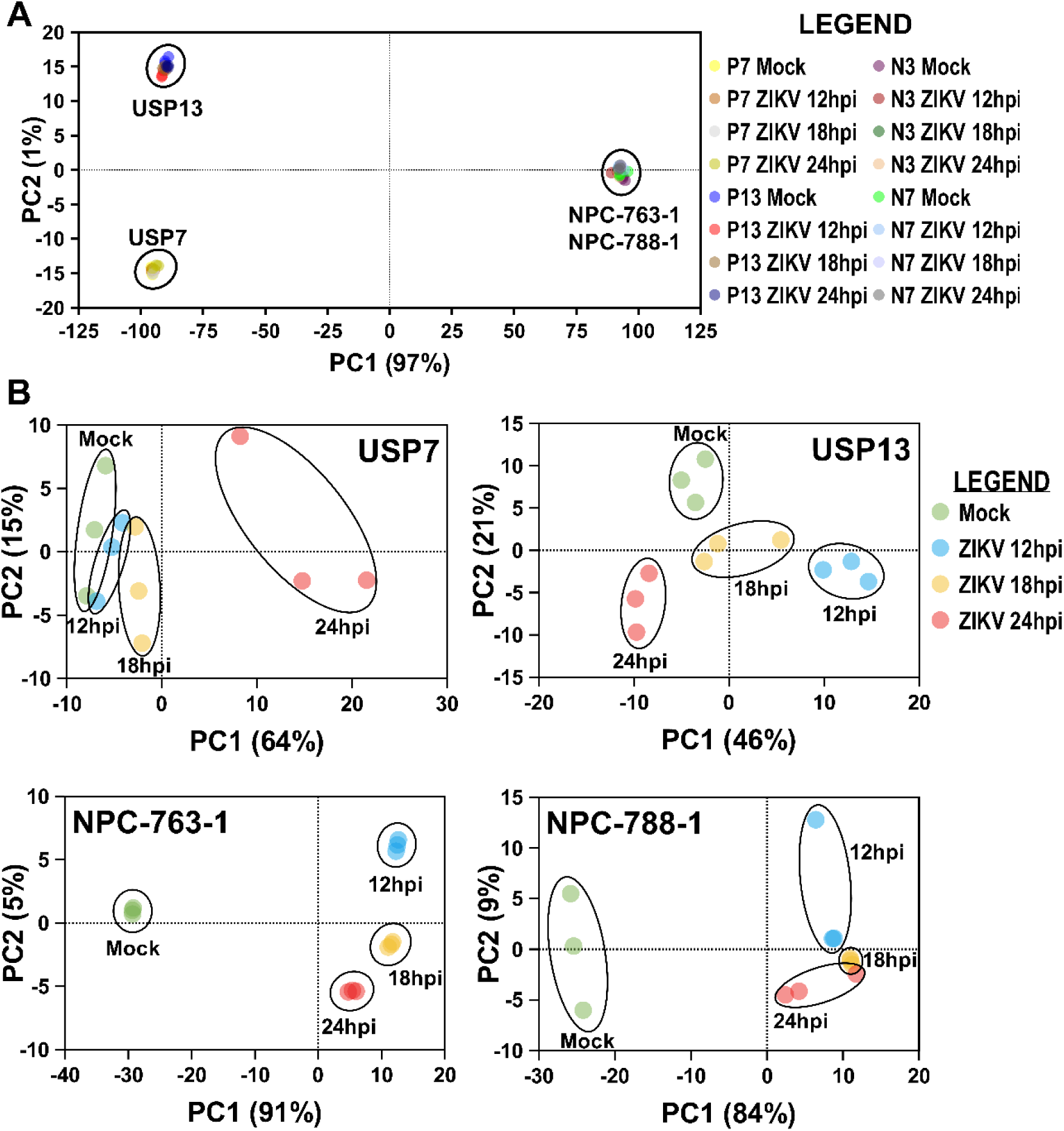
PCA of ZIKV-infected RNA-Seq samples. PCA plots of **(A)** all RNA-Seq samples together and **(B)** individual cell lines. Abbreviations, Zika virus (ZIKV), USP7 (P7), USP13 (P13), NPC-763-1 (N3), NPC-788-1 (N7), hours post-infection (hpi), Principal Component (PC).

